# Single-cell transcriptomic dynamics exposes hidden survival trajectory under antibiotic treatment

**DOI:** 10.64898/2025.12.03.692042

**Authors:** Jinxin Zhao, Ning Liu, Xingjian Wang, Jiru Han, Yu-Wei Lin, Panjie Hu, Heidi H. Yu, Hasini Wickremasinghe, Jing Lu, Yan Zhu, Xiaoting Hua, Yunsong Yu, Hua Wu, Jianzhong Ye, Tieli Zhou, Tony Velkov, Jian Li

**Affiliations:** Infection Program and Department of Microbiology, Monash Biomedicine Discovery Institute, Monash University, Clayton, VIC 3800, Australia; Monash University - Wenzhou Medical University Biomedical Alliance, Clayton, VIC 3800, Australia; Consortium for Infection and Innovation (CII), Monash University - Wenzhou Medical University Biomedical Alliance, The First Affiliated Hospital of Wenzhou Medical University, Wenzhou, Zhejiang, China; Centre of Excellence for Antimicrobial Therapeutics Discovery and Innovation, Monash Suzhou Research Institute, Suzhou Campus, Jiangsu 215123, China; South Australian immunoGENomics Cancer Institute (SAiGENCI), Faculty of Health and Medical Sciences, The University of Adelaide, Adelaide, SA 5005, Australia; Adelaide Centre for Epigenetics, School of Biomedicine, Faculty of Health and Medical Sciences, The University of Adelaide, SA 5005, Australia; Genetics and Gene Regulation Division, The Walter and Eliza Hall Institute of Medical Research, Parkville, VIC 3052, Australia; Key Laboratory of Clinical Laboratory Diagnosis and Translational Research of Zhejiang Province, Department of Clinical Laboratory, The First Affiliated Hospital of Wenzhou Medical University, Zhejiang, China; Institute of Infectious Disease, The Second Affiliated Hospital of Tianjin Medical University, Tianjin, China; Systems Biology Centre, Tianjin Institute of Industrial Biotechnology, Chinese Academy of Sciences, Tianjin, China; Department of Infectious Diseases, Sir Run Run Shaw Hospital, Zhejiang University School of Medicine, Zhejiang, 310001, China; Department of Pharmacology and Toxicology, College of Veterinary Medicine, Henan Agricultural University, Henan, China; Department of Pharmacology, Monash Biomedicine Discovery Institute, Monash University, Clayton VIC 3800, Australia

**Keywords:** Antimicrobial resistance, *Acinetobacter baumannii*, heterogeneity, polymyxins, single-cell transcriptomics

## Abstract

Heterogeneity plays a major role in bacterial resistance to antibiotic treatment while the mechanism at single-cell level is largely unknown. Here, we employed a robust integration of bulk and bacterial single-cell RNA-seq (scRNA-seq) to uncover how individual cells of *Acinetobacter baumannii* reorganise the heterogenous transcriptome in response to antibiotic. Using polymyxin as a representative, bulk RNA-seq showed canonical envelope- and efflux-centred responses but obscured underlying heterogeneity. Single-cell profiling resolved these averages into discrete subpopulations whose abundances shifted with concentration and time. Specifically, in early time an envelope-stress programme predominated survival at the low concentration, whereas an outer-membrane repair/efflux programme dominated survival at the high concentration. These patterns revealed structured and time-resolved heterogeneity, highlighting an ingenious bacterial stress responsive strategy. Through trajectory inference, we further revealed a concentration-dependent bifurcation of cell fates: low-concentration treated cells detoured through a transient stress state and rejoined growth, whereas high-concentration treated survivors diverted into a slow-growing, tolerant branch. Perturbing marker genes from these programmes altered fitness eventually and rapidly increased permeability, depolarisation and reactive-oxygen burden, linking state to survival capability. Collectively, we map a dynamic, concentration-structured landscape of antibiotic responses at the single-cell level, revealing diverse survival trajectories that are obscured in conventional population-averaged analyses. Our developed single-cell based framework revealed tolerant bacterial cells with rewiring of the transcriptional landscape, causing emergence of antibiotic resistance. Importantly, these findings urge precision antimicrobial therapy in patients to minimise emergence of antibiotic resistance.

## Main

Current microbiology remains largely grounded in population-level approaches and bulk transcriptomic analyses based on bacterial populations have significantly advanced our understanding of bacterial adaptability to antibiotic pressure^1–4^. For several decades, these population-averaged studies have defined the core principles of antibiotic resistance. For example, in *Acinetobacter baumannii*, a top-priority pathogen identified by the World Health Organization (WHO), exposure to the last-resort polymyxins induces modifications of lipid A in lipopolysaccharide, upregulates efflux systems, and triggers protective envelope responses^5,6^.

Yet bulk approaches report only averaged gene expression of millions of cells, obscuring critical heterogenous phenotypic differences between individual bacterial cells^7,8^. Bacterial populations commonly adopt bet-hedging strategies in which a minority of cells enter protective, low-activity states while other cells follow different programmes, ensuring survival under stress^9,10^. In polymyxin-heteroresistant *A. baumannii*, a subpopulation of cells can survive high antibiotic concentrations of polymyxin despite the overall population being susceptible^11^. Rare survivors revert to growth phase once the antibiotic stress is removed, leading to infection relapse^12–14^. Pinpointing these defence programmes to specific cells and tracking their trajectories across concentration and time remains a critical gap in antimicrobial resistance research^15^.

Here, we integrate both population- and single-cell transcriptomics to map the concentration-and time-dependent adaptive landscape of 72,176 individual cells of *A. baumannii* following antibiotic exposure (**Fig. 1A**), using polymyxin as an exemplar as it serves as a last-line therapy against multidrug-resistant (MDR) Gram-negative infections^16^. We reasoned that transcriptional heterogeneity within tolerant subpopulations is obscured by population averages. Bulk RNA-seq shows recover canonical enveloped-and efflux-centred stress responses, whereas scRNA-seq uncovers how these programmes are partitioned among subpopulations and reweighted with concentration and time. Perturbating state-defining genes functionally connects these programmes to quantitative susceptibility phenotypes, uncovering tractable antimicrobial vulnerabilities. Together, our integrated single-cell approach resolves population averages into discrete cell state and reveals an early, concentration-dependent bifurcation in fate. These genetic and physiological concordances map transcriptional state onto tolerance and survival, and define a framework for establishing precision therapies against life-threatening infections in patients by eradicating resilient antibiotic-resistant subpopulations.

**Figure 1:**
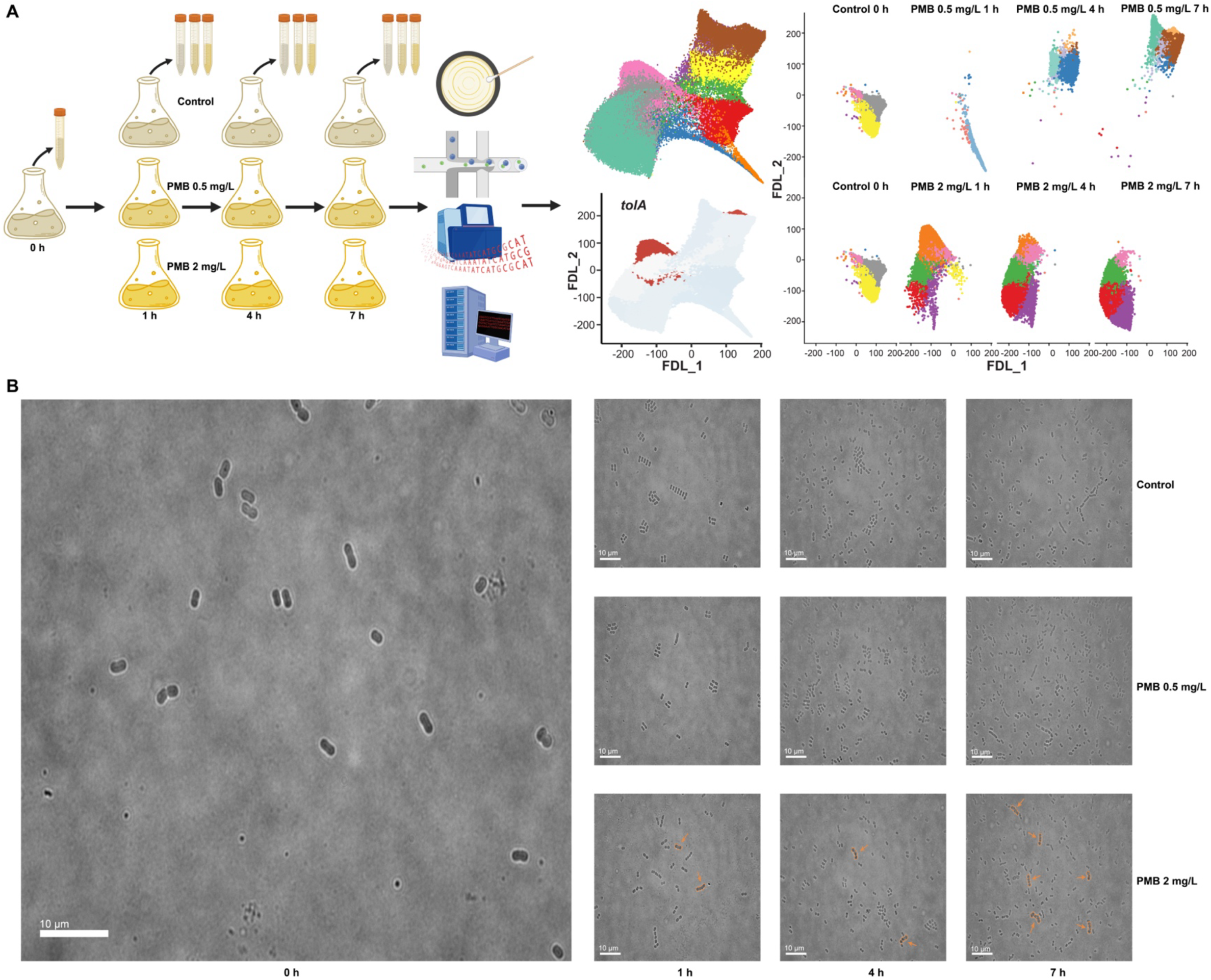
Study design and time-lapse imaging of *A. baumannii* under polymyxin treatment. **(A)** Schematic of the experimental design for single-cell RNA-seq. *A. baumannii* strain AB5075 cultures were grown to mid-log phase and then exposed to polymyxin B at either 0 mg/L (control), 0.5 mg/L (low), or 2 mg/L (high) based on its pharmacokinetics in patients. Samples for single-cell RNA-seq were collected at 0, 1, 4, and 7 h after antibiotic addition. Cells were immediately fixed to stabilise RNA profiles, then processed with the smRandom-seq droplet platform to capture and barcode single-cell transcriptomes, followed by sequencing and computational analysis. **(B)** Time-lapse microscopy stills of *A. baumannii* under the same conditions, taken at 0 h, 1 h, 4 h, and 7 h. The images illustrate growth dynamics and morphological changes: in the control, cells grow and divide normally; under 0.5 mg/L polymyxin B, minor growth occurs; under 2 mg/L, rapid killing occurs, and a small subset survives is slowed with some cell chaining or filamentation (arrowheads indicate cells forming clusters). These observations motivated downstream single-cell transcriptomic analysis.

## Results

### Bulk transcriptomics masks heterogeneous bacterial responses to antibiotic treatment

We first established the population-level transcriptional response of *A. baumannii* (strain AB5075) to polymyxins using bulk RNA-seq. Mid-log phase cultures were exposed to low (0.5 mg/L, 1× MIC) or high (2 mg/L, 4× MIC) concentrations based on polymyxin pharmacokinetics in patients^17^. As expected, polymyxin elicited a rapid reprogramming of the bacterial transcriptome, dependent on antibiotic concentration and exposure duration. Principal component analysis (PCA) showed clear segregation of samples by treatment and timepoint, indicating that polymyxin treatment rapidly shifts the global gene expression away from the untreated control even after 1 h (**Fig. 2A**). Notably, low and high concentrations separated along distinct principal component axes, showing quantitative and qualitative differences in their transcriptomic responses.

**Figure 2:**
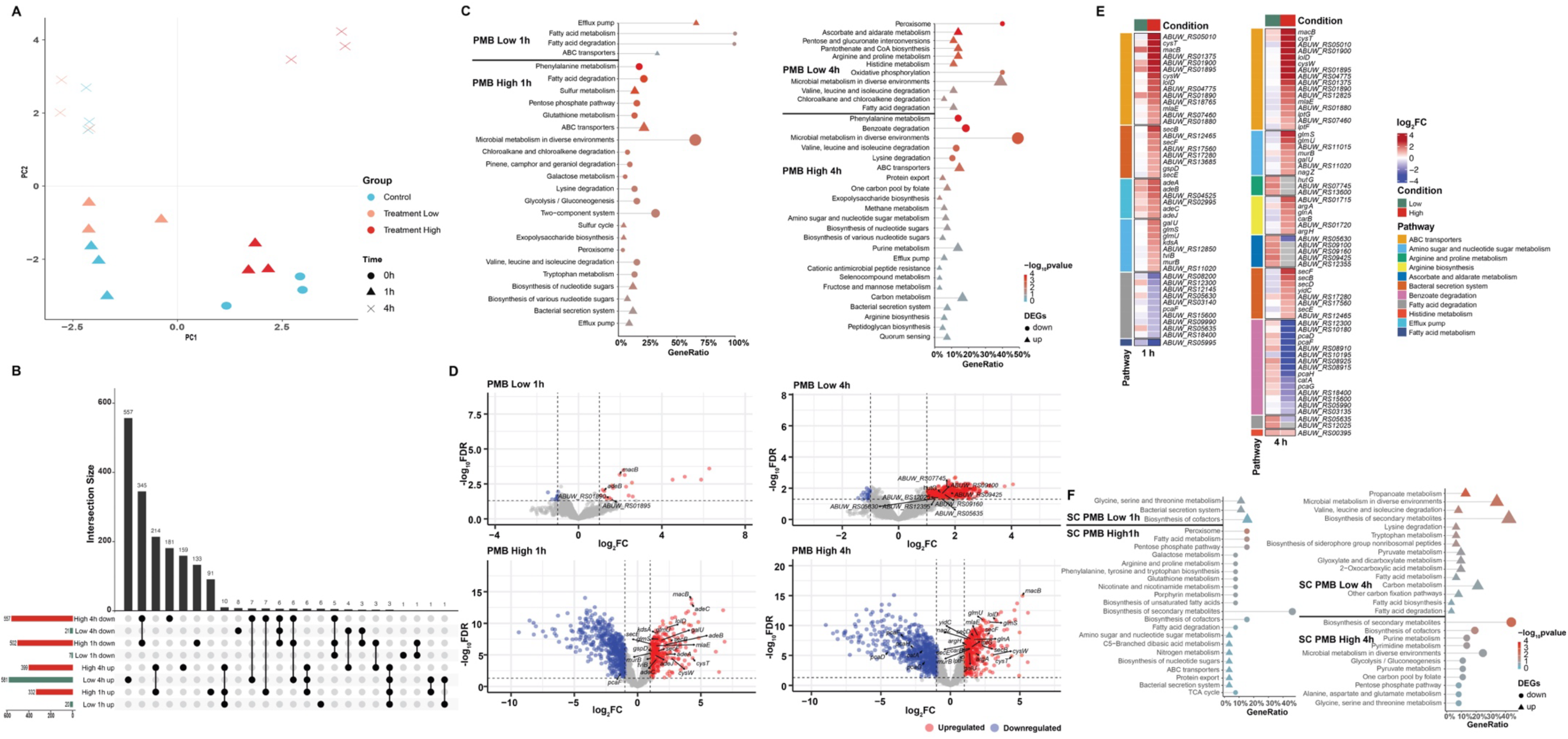
Bulk transcriptomic analysis of *A. baumannii* under polymyxin B (PMB) and comparison to the data from single-cell approach. **(A)** PCA of bulk RNA-seq profiles under polymyxin B treatment (Low: 0.5 mg/L; High: 2 mg/L) at 1 h and 4 h, versus untreated controls (0 h). Points represent individual samples (biological replicates), coloured by polymyxin concentration and shaped by time. PC1 and PC2 capture the major variance, separating conditions by antibiotic exposure and highlighting differences between low and high concentrations. **(B)** UpSet plot of DEGs across four pairwise comparisons, revealing a core stress response as well as unique subsets induced by high concentration or timepoints. **(C)** KEGG pathway enrichment of up-regulated DEGs per condition. The top enriched pathways are shown for each condition (dot size corresponds to gene ratio within the pathway, and colour indicates statistical significance). Key pathways include efflux pump systems (e.g., *ade* operon), and ABC transporters, and various metabolic and stress response pathways, aligning with known mechanisms of polymyxin action and resistance. **(D)** Volcano plots of differential expression results at 1 h (top) and 4 h (bottom) for Low (left) and High (right) concentrations. Each point is a gene; red and blue points denote significantly up- and down-regulated genes, respectively (FDR < 0.05). The high concentration triggers a broader response by 4 h, including pronounced down-regulation of housekeeping genes and strong up-regulation of stress adaptive genes, whereas the low concentration causes a milder shift. **(E)** Pathway signatures of the polymyxin response. Heat maps show differentially expressed genes (FDR < 0.05) grouped by functional modules under low and high polymyxin concentrations at 1 h and 4 h. Early responses centre on envelope and efflux activation, while later stages show sustained membrane repair and metabolic adaptation. **(F)** KEGG pathway enrichment analysis of genes identified through single-cell RNA-seq as differentially expressed in specific subpopulations under polymyxin B (at 1 h and 4 h, for low and high concentrations).

Differential expression analysis revealed 27-1,000 significantly regulated DE genes per condition. The low concentration caused a relatively modest transcriptional perturbation at early time points (only 27 differentially expressed genes (DEGs) at 1 h), which expanded to a few hundred by 4 h as the stress persisted (**Fig. 2B**). In contrast, the high concentration triggered much broader responses immediately, over 800 DEGs by 1 h, with much more pronounced differences by 4 h (**Fig. 2B-D**). It is evident that *A. baumannii* mounts a graded response to polymyxin, with the higher concentration activating more stress pathways on top of the core response due to the lower stress level.

Pathway enrichment analyses underscored major stress responses (**Fig. 2C**). Both concentrations induced a common set of early defense genes, notably efflux and transporter systems, like *adeABC* and *macAB*-*tolC*, which were upregulated within 1 h (**Fig. 2B–D**). However, high-concentration treatment uniquely drove strong upregulation of envelope biogenesis and repair pathways, consistent with more severe membrane damage. Key induced genes included the *mla* retrograde phospholipid transport system (*mlaF, mlaD, mlaC* and *mlaB*), lipopolysaccharide transports (*lptG* and *lptF*), and outer-membrane lipoprotein trafficking factors (*lolD*, *glmS*, *glmU*) (**Fig. 2D**, **E** **and Extended Data Fig. 1 and Fig. 2**). These systems maintain membrane integrity and are essential for *A. baumannii* survival during polymyxin treatment. Similarly, genes encoding protein secretion and membrane insertion machineries (*secB*, *secD-F*, *gspC*, *gspD*, *gspO* and *YidC*) were upregulated under the high-concentration treatment (**Fig. 2E, Extended Data Fig. 1 and Fig.2**), consistent with an increased demand for protein export and membrane remodelling during polymyxin-induced damage. Beyond membrane-focused defense, high-concentration polymyxin also caused marked shifts in cellular metabolism. Notably, the arginine biosynthesis and catabolism genes (e.g., *argA*, *carB*, *argH* and *glnA*) were strongly upregulated (**Extended Data Fig. 1**), suggesting a protective role for arginine metabolism in membrane stabilisation, oxidative stress mitigation, and generating antioxidant precursors like polyamines. Conversely, the high concentration selectively repressed genes for degrading aromatic compound (the *pca* operon) (**Extended Data Fig. 1**), presumably to conserve resources for stress survival. Likewise, ascorbate and aldarate metabolism genes were differentially expressed (**Fig. 2E**), aligning with their potential role in metabolic flexibility and antioxidant defense, though their precise contribution to polymyxin survival remains to be clarified.

Together, these bulk RNA-seq data depict a concentration-dependent stress program in *A. baumannii*, mainly efflux pumps, transporters, envelope lipid recycling, secretion/insertion systems, oxidative stress defense, and metabolic rerouting. Importantly, the breadth and concentration-specificity of these changes hint that not all bacterial cells in the population contribute equally to the responses, raising the possibility of hidden heterogeneity that cannot be resolved by bulk analysis. We therefore performed scRNA-seq to dissect how these transcriptional changes are partitioned across individual cells (**Fig. 2F**).

### Single-cell RNA sequencing delineates heterogeneous subpopulations in *A. baumannii*

To resolve this heterogeneity in *A. baumannii* (**Fig. 1B**), we employed scRNA-seq mirroring the bulk RNA-seq experiment (control, 0.5 mg/L [low] or 2 mg/L [high] polymyxin B at 0, 1, 4, and 7 h). Utilising smRandom-seq, we encapsulated and processed 1,000,000 individual bacteria from these samples and obtained high-quality transcriptomes for ~80,000 single cells after filtering. On average approximately 200 genes were detected per cell (**Extended Data Fig. 3**), providing sufficient depth to resolve distinct cell states, including those with modest gene expression changes. Pathway enrichment from single-cell DEGs of 72,176 single cells recapitulated mirrored bulk signatures at the population level (**Fig. 2F** vs **2C**), converging on ABC transporters, amino acid metabolism, and envelope maintenance as central polymyxin stress responses. This concordance validated the robustness of the single-cell dataset while enabling resolution of these responses at the cellular scale.

Surprisingly, unsupervised clustering identified 15 transcriptionally distinct subpopulations (clusters) (**Fig. 3A, B**). Each subpopulation also comprised a mixture of clusters with different transcriptomic states, and the proportions of these states shifted systematically with concentration and time (**Fig. 3A, B and Extended Data Fig. 4**). We visualised the relationships between cells using a force-directed layout (FDL) graph of the *k*-nearest-neighbour cell network. In this two-dimensional projection, cells with similar transcriptomes clustered together into discrete groups corresponding to the 15 identified clusters (**Fig. 3A, B** and **Extended Data Fig. 4**). Cells from different conditions frequently co-localised in this embedding, revealing baseline heterogeneity even in the absence of drug, and antibiotic exposure then reweighted this baseline towards specific states.

**Figure 3:**
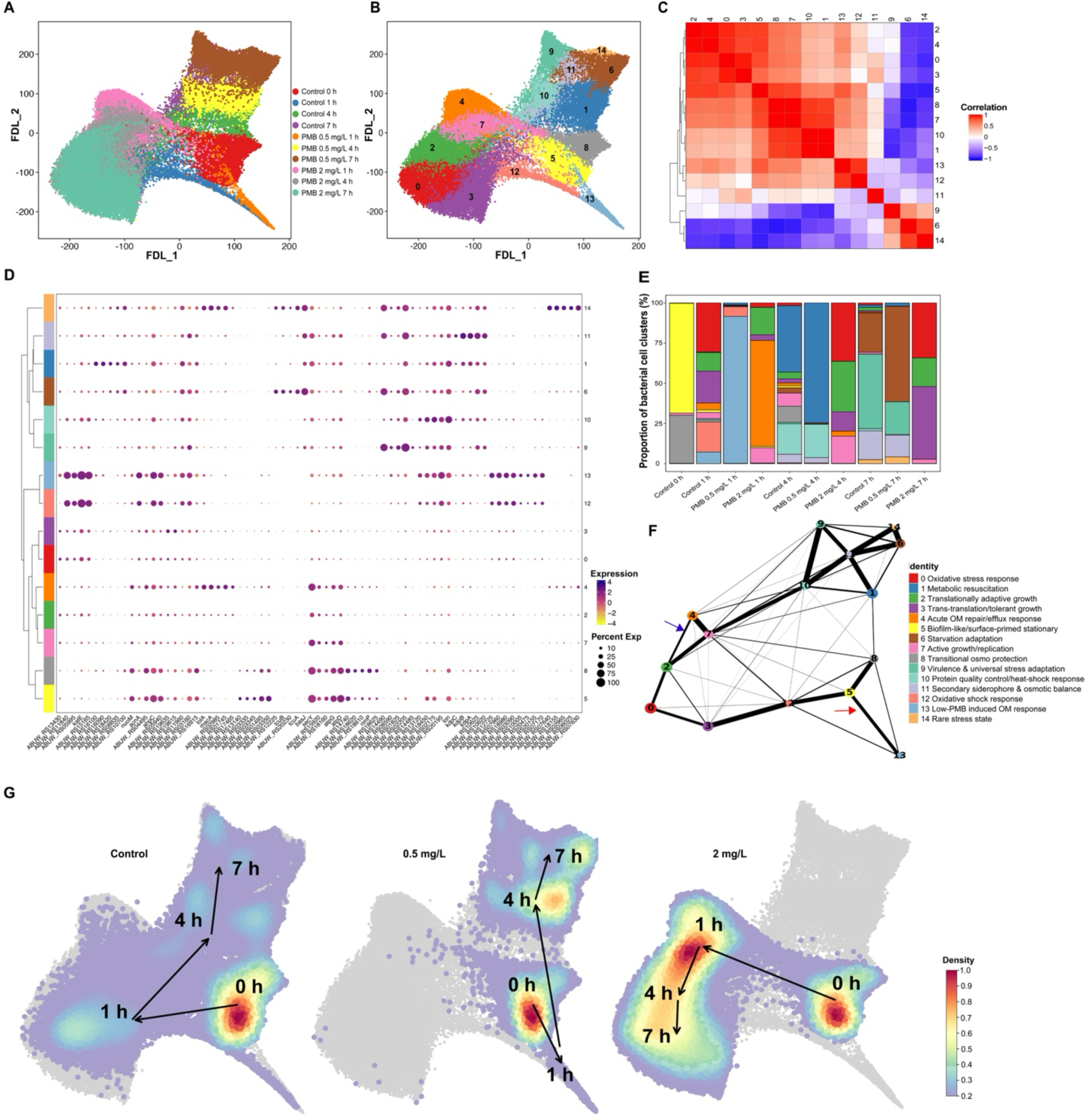
Single-cell transcriptomics uncovers extensive heterogeneity in the polymyxin response of *A. baumannii*. **(A)** Force-directed layout (FDL) projection of all sequenced single cells coloured by treatment group: control (no antibiotic), low-concentration polymyxin B (PMB, 0.5 mg/L), and high-concentration PMB (2 mg/L) at 0 h, 1 h, 4 h, and 7 h. Each point represents the transcriptome of a single bacterial cell; points clustering together indicate similar gene expression. The overlap of colours in several regions illustrates that cells from different conditions can occupy similar transcriptional states, but the relative abundance of those states differs by condition and time. **(B)** FDL projection (same as in A) coloured by cluster identity (15 distinct clusters, numbered 0–14). This view delineates the transcriptionally distinct subpopulations present in the combined dataset. Each cluster corresponds to a unique gene expression program (see panel D and Extended Data Table 1 for functional characterisation). **(C)** Transcriptional relatedness of single-cell states. Heat map showing pairwise Pearson correlations between cluster-centroid expression profiles (computed from the variable genes used for clustering). Rows/columns are ordered by hierarchical clustering. The block structure groups biologically related programmes and this correlation structure underlies the trajectory bifurcation shown in panel **F**. **(D)** Dot plot of the mean expression of select marker genes (rows) in each cluster (columns). Dot colour intensity represents the average expression level of the gene within that cluster, and dot size corresponds to the fraction of cells in the cluster expressing the gene. For each cluster, five marker genes (by the |Log_2_FC| rank) are selected to illustrate the distinct molecular signatures underlying each subpopulation. For example, cluster 4 shows high expression of *macB* (efflux pump) and *lolA*, cluster 13 is characterised by *nlpE* and *ymgE*. These markers underpin the functional annotations of clusters (e.g., envelope stress). **(E)** Stacked bar plots showing the proportion of cells belonging to each cluster in each condition over time (control vs 0.5 mg/L vs 2 mg/L at 0, 1, 4, 7 h). This highlights how the population composition shifts: such as cluster 13 (purple) dominates the 1-h low-concentration sample, whereas cluster 4 (green) dominates the 1-h high-concentration sample, and control 1-h cells are distributed among clusters 0, 2, 3, etc. By 7 h, low-concentration and control have a more similar distribution (mostly clusters 6, 9, 11 for stationary phase), whereas high-concentration remains distinct. **(F)** FDL graph with partition-based graph abstraction (PAGA) of the single-cell data overlay, summarising lineage relationships between clusters. This reinforces the branching pattern of cell-state transitions under polymyxin stress. Each node is a cluster (numbered, with size roughly proportional to cluster frequency) and edges connect clusters that have transcriptional continuity (weighted by inferred transition probability). The layout highlights a bifurcation: clusters associated predominantly with high-concentration treatment of polymyxin form a separate branch (right side, including clusters 4 → 2 → 0/3) apart from the main growth-and-stationary-phase branch (left side, including clusters 5/8 → 0/12/13 → 1/7/10 → 6/9/11). Together, panels A–F show that scRNA-seq reveals a complex, dynamic mosaic of *A. baumannii* phenotypes under polymyxin exposure, ranging from vulnerable, dying cells to adaptive survivors, an intricacy that traditional bulk RNA-seq cannot resolve. (**G**) Manifold-based density maps and population-flow trajectories reveal concentration-dependent rewiring of the transcriptional landscape. Single-cell transcriptomes were projected into the shared FDL manifold (grey). Two-dimensional kernel density estimates were computed for cells from each trajectory and normalised to a common scale (colour bar). Arrows connect the centroids of population distributions across time (0 h → 1 h → 4 h → 7 h). Control cells follow the normal growth trajectory, cells under low-concentration treatment briefly diverge into an envelope-stress region before rejoining the main path, while cells under high-concentration treatment enter a distinct acute-stress/dormancy region from which they do not return. Together, these density and flow maps depict the geometric divergence of antibiotic stress responses over time.

We next investigated the marker gene signatures of each cluster to assess their identities and biological significance. We observed that each cluster had a unique transcriptional profile indicative of a specific adaptive strategy and physiological status, spanning growth and resuscitation, oxidative-stress tolerance, osmoadaptation and envelope-centred defences (**Fig. 3C-D** and **Extended Data Table 1**). These 15 identified clusters delineate a rich landscape of *A. baumannii* cell states during treatment, underscoring the heterogeneity that was masked in the bulk averages (**Fig. 3C-E** and **Extended Data Table 1**). For example, in both untreated and polymyxin-treated samples, growth-associated and stress-associated subpopulations coexisted (**Fig. 3A, B** and **E**). However, the balance between these cell states differed markedly depending on the presence and concentration of polymyxin (**Fig. 3E**). Under the low concentration, the population was dominated by a single subpopulation: by 1 h, ~91% cells belonged to cluster 13. This cluster represents an envelope-stress response state (mainly upregulating *nlpE*-family and other envelope stress factors) that was only a minor fraction of the untreated control. In contrast, under the high concentration, a different subpopulation (cluster 4) became predominant, mounting an intense efflux and membrane repair response (e.g., upregulating *macAB*-*tolC*, *lolA*) (**Fig. 2B, 2D** and **Extended Data Fig. 5**). In the same high-concentration culture, other cells simultaneously occupied very different states, some remained in a growth-associated cluster (cluster 7), while others adopted slow-growing, stress-tolerant phenotypes akin to stationary phase (e.g., cluster 5) (**Extended Data Fig. 4**). Thus, the population-level response to antibiotic treatment is a composite of disparate single-cell behaviours. Bulk RNA-seq could only capture the average effect of the antibiotic, whereas the single-cell approach unmasks how the stress responses are partitioned across diverse subpopulations and individual cells. In summary, our cluster analysis demonstrates that polymyxin-treated *A. baumannii* cultures consist of a mosaic of coexisting cell states, each deploying a distinct transcriptional defense strategy rather than mounting a uniform, population-wide stress response as shown by the bulk RNA-seq data.

### Antibiotic treatment reshapes the transcriptional landscape, enriching distinct subpopulations

Polymyxin exposure fundamentally reshaped the distribution of transcriptional states in the population by preferential expansion of certain subpopulations. We firstly established the baseline dynamics without polymyxin. Even at time 0, cells were heterogenous: the inoculum of ~1 million cells from an overnight stationary-phase culture consisted mainly of stationary phase-like states (clusters 5 and 8) (**Fig. 2E, Extended Data Fig. 4**). Upon transfer to fresh medium (untreated control), the population progressed through a predictable succession as it grew. By 1 h, many control cells had exited stationary phase and entered rapid growth or early stress response clusters (0, 2, 3, 7, 12, 13) (**Fig. 3E**). By 4 h (mid-log phase), as nutrients were consumed, the population shifted toward metabolic resuscitation (cluster 1, likely reflecting awakening from dormancy) and proteostasis management (cluster 10). By 7 h, the control population predominantly occupied clusters 6, 9, and 11, indicating adaptation to nutrient limitation (e.g., increased iron scavenging) and stationary phase survival (mimicking late stationary phase or biofilm-like state). This temporal pattern reflects the intrinsic growth cycle progression of *A. baumannii* subpopulations, providing a baseline trajectory for comparison (**Fig. 3D, E**, **Extended Data Fig. 4** and **Table 1**).

Polymyxin treatment dramatically redirected the population into distinct transcriptional states rarely observed in the drug-free condition. At low concentration, cells converged by 1 h on an envelope-stress programme (cluster 13) centred on periplasmic buffering and osmoprotection, marked by GlsB/YeaQ/YmgE- and NirD/YgiW/YdeI-family stress factors, trehalose biosynthesis (*otsB*), small chaperones and outer-membrane lipoproteins (**Extended Data Fig. 5A**). Among cluster 13 markers, *ABUW_RS18980* (GlsB/YeaQ/YmgE-family stress response membrane protein) and *ABUW_RS07170* (NirD/YgiW/YdeI-family stress tolerance protein) typify envelope-stress mitigation stabilising the periplasm and cell surface under mild perturbation. Induction of *otsB* (trehalose-phosphate phosphatase) indicates flux through trehalose biosynthesis, an osmoprotective route that preserves membrane integrity and protein function^18^. A hemerythrin-domain protein (*ABUW_RS01615*) and a small heat-shock chaperone (*ABUW_RS07965*) point to redox control and proteostasis. Enrichment of lipoproteins (*ABUW_RS20155*, *ABUW_RS20475*) together with DUF-annotated periplasmic proteins suggests reinforcement of outer-membrane interfaces. The coherence of these marker genes supports a rapid, synchronised envelope-stress adaptation across most cells at low concentration. This response was relatively transient. By 4 h, the low-concentration population largely realigned with the control trajectory, transitioning into growth recovery and protein homeostasis (cluster 1 for metabolic resuscitation, cluster 10 for proteostasis stress), similar to the control at 4 h. By 7 h, the low-concentration culture settled primarily into stationary-phase clusters (6, 9, 11), akin to drug-free control but with a subtle shift toward stronger iron-scavenging and nutrient-starvation signatures, perhaps due to combined polymyxin stress and nutrient depletion. In summary, low-concentration polymyxin triggers a buffer-and-recover module in which cells transiently fortify the envelope, attenuate nascent oxidative and proteotoxic stress, and then re-enter a growth trajectory comparable to normal growth (**Fig. 3D, E** and **Extended Data Fig. 4**).

The high-concentration treatment enriched cluster 4, an envelope-repair and efflux pump module for acute envelope injury (**Fig. 3E**). Prominent markers include the MacAB-TolC ABC transporter and the AdeABC RND system, together consistent with immediate antibiotic extrusion of polymyxin and removal of damaged lipids (**Extended Data Fig. 4**). The lipoprotein chaperone *lolA* denotes active lipoprotein trafficking to the outer membrane, a prerequisite for rebuilding the envelope under stress. The two-component system BfmS/BfmR appears alongside an OmpA-like glycine-zipper membrane protein (*ABUW_RS07630*), consistent with envelope-damage sensing and porin remodelling. Ancillary factors, including an RcnB-family protein (*ABUW_RS18750*) and SIMPL-domain proteins, support metal and envelope homeostasis during repair (**Extended Data Fig. 4**). Cells in this cluster thus prioritise their transcriptional responses in lipoprotein delivery, membrane rebuilding and multi-systems efflux pump rather than in unperturbed growth (**Fig. 3E**, **Extended Data Figs. 5B and 6**). Notably, cluster 4 is rare at low concentration, implying this intense repair/efflux response is selectively recruited under higher antibiotic burden. At later timepoints, high-concentration populations did not revert to exponential-growth clusters seen in the recovering low-concentration culture. Instead, by 4 h, the composition shifted toward oxidative stress tolerance (cluster 0), attenuated yet active growth (cluster 2), and translational recovery/tolerant growth state (cluster 3) (**Extended Data Fig. 4**). By 7 h, surviving cells predominantly occupied clusters 0 and 3 (**Extended Data Fig. 4**), maintaining stress-mitigation systems (e.g., antioxidants in cluster 0), active ribosome quality-control via trans-translation (cluster 3), and drastically curtailed growth-related activities. This high-concentration scenario reflects selection and amplification of a tolerant subpopulation that sacrifices growth for survival, aligning with viability measurements. The high-concentration treatment caused an ~99% drop in bacterial number by 4 h, yet a small fraction of cells survived and began regrowing by 7 h (**Fig. 1B** and **Extended Data Fig. 6**). In contrast, the low concentration caused only a transient 1–4 h growth delay with most cells remaining viable, consistent with transcriptional recovery evidence.

Subsequently, we conducted trajectory analysis to show the transcriptomic dynamics at the single-cell level, further emphasising these divergent paths (**Fig. 3E-G**). In the absence of antibiotic, cells followed a linear trajectory through lag, exponential, and stationary phase clusters. Under the low-concentration treatment, the trajectory initially split off at cluster 13 (reflecting 1 h envelope-stress detour) but then merged back into the main stationary-phase progression (clusters 6, 9, 11) by 7 h (**Fig. 3E-G** and **Extended Data Fig. 4A**). In contrast, the high-concentration condition formed a separate branch: after cluster 4 at 1 h, the graph veered onto a path connecting clusters 4 → 2 → 0/3, distinct from control or low-concentration routes (**Fig. 3F** and **G**).

Thus, the trajectory map highlights two distinct divergent polymyxin stress outcomes. One branch (low-concentration treatment) leads to clusters associated with adaptive stress responses followed by regrowth, rejoining the normal trajectory after the initial detour (**Fig. 3F blue arrow** and **3G**). The other branch (high-concentration treatment) leads to deep stress and dormancy, culminating in tolerant clusters with no regrowth during the experiment (**Fig. 3F red arrow** and **3G**). These distinct dynamics illustrate how polymyxin fundamentally reprograms the population’s lineage structure, promoting specialised survival strategies in a subset while the rest perish. Essentially, the single-cell data capture a phenotypic bifurcation, where one subpopulation actively mounts defenses and survives in a slowed, protective state, while another subpopulation fails to adapt and is eliminated. Notably, the surviving branch corresponds to cells that, through reversible phenotypic changes, tolerate the antibiotic long enough to eventually resume growth once the stress abates.

### Functional relevance of subpopulation-specific gene expression programs

Trajectory analysis placed an envelope-stress programme at low concentration (cluster 13, periplasmic protectants, osmolyte synthesis and chaperones) and an acute outer-membrane repair and efflux response programme at high concentration (cluster 4, lipoprotein sorting, efflux pump and envelope-damage signalling). We therefore investigated whether genes marking these dominant single-cell states are mechanistically linked to survival under antibiotic exposure. We selected marker genes spanning clusters 4 and 13 at 1 h and profiled the corresponding *A. baumannii* AB5075 transposon mutants; particularly, we (i) quantified 24-h fitness (WT-normalised area-under-the-curve (AUC)) across control, 0.5 mg/L, and 2 mg/L polymyxin B (**Fig. 4A** and **Extended Data Fig. 7)**, and (ii) measured 1-h physiological readouts of envelope permeability (PI), membrane depolarisation DiBAC_4_(3) depolarisation, and CellROX fluorescence under the same treatments to assess envelope integrity, membrane potential, and oxidative stress (**Fig. 4B–D**). Across the panel, 24-h fitness shifted relative to the wild-type in a concentration-dependent manner, with many mutants showing the largest reductions in AUC at 2 mg/L (**Fig. 4A**). Representative growth curves (**Extended Data Fig. 7C**) visually highlight this trend: under 2 mg/L, mutants in OM repair/efflux (e.g., *macB*, *adeA, tolA*) and envelope-stress signalling (e.g., *bfmS*, *ABUW_RS07170*) exhibited prolonged lag, slower growth, and lower final optical densities, whereas trajectories in the control and at 0.5 mg/L were near the wild-type or only modestly delayed.

**Figure 4:**
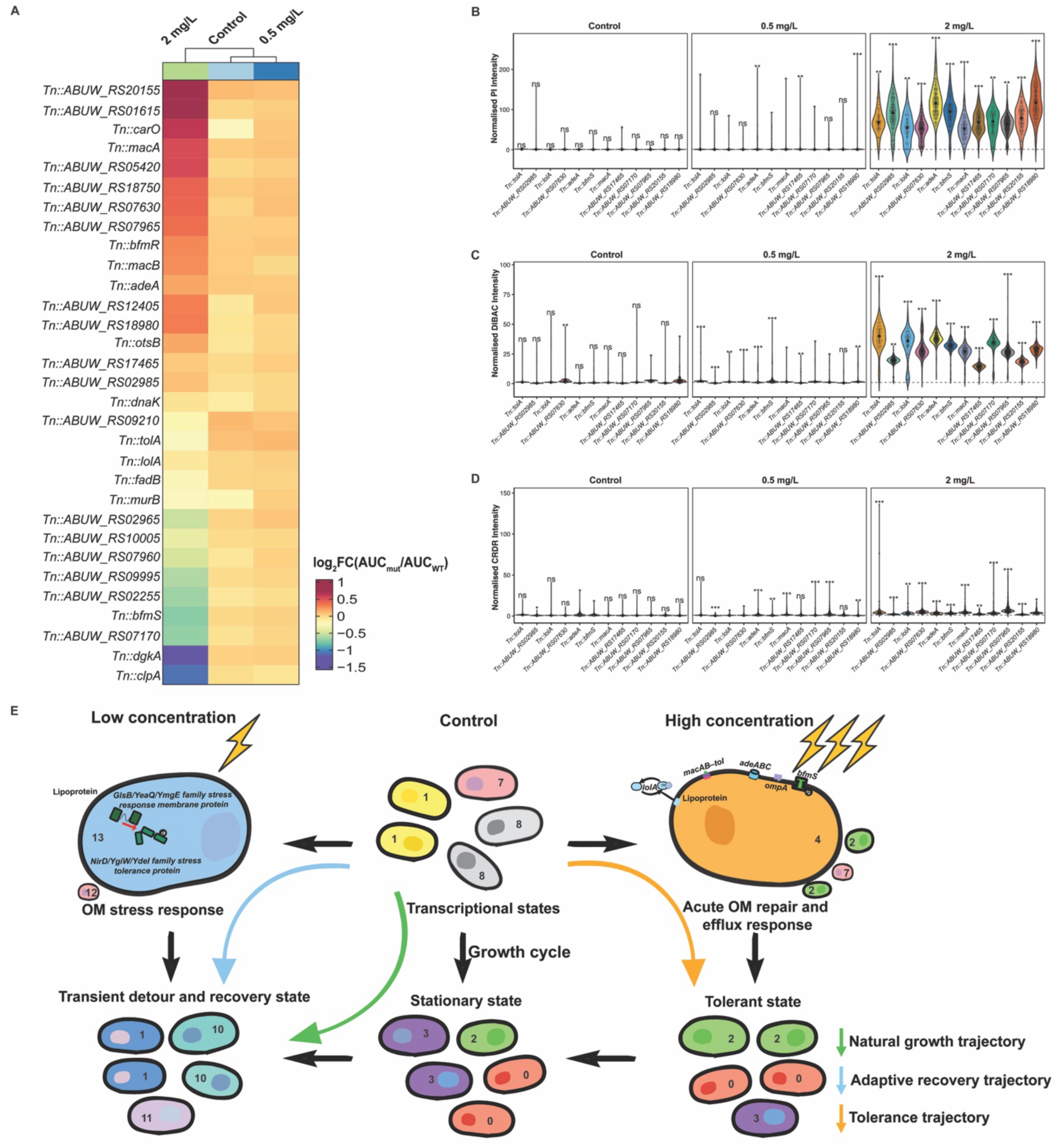
Single-cell defined subpopulation programs nominate genes that determine survival under membrane-targeting antibiotic stress. **(A)** Fitness landscape of marker-gene mutants across concentration treatments. Heat map of 24-h growth fitness for AB5075 transposon mutants in genes that mark the dominant single-cell subpopulations at 1 h (OM repair/efflux and envelope-stress programs). Values are log_2_FC(AUC_mut_/AUC_WT_) from optical-density trajectories, normalised within condition; more negative values indicate greater growth impairment. **(B–D)** Orthogonal single-cell readouts reveal the physiological basis of susceptibility at 1 h. Violin plots show WT-normalised fluorescence distributions for **(B)** propidium iodide (PI; envelope permeability), **(C)** DiBAC_4_(3) (membrane depolarisation), and **(D)** CRDR (oxidative stress) measured in the same mutant panel under control, 0.5 and 2 mg/L polymyxin B treatments. At high concentration, mutants in OM maintenance/efflux and stress signalling (e.g., *tolA*, *lolA*, *adeA*, *bfmS*, *ABUW_RS07170*) exhibit increased PI and DiBAC signals and elevated CRDR, consistent with a permeable, depolarised, ROS-burdened state. At low concentration, distributions are largely indistinguishable from WT at the 1-h snapshot, consistent with a transient envelope-stress state inferred from the single-cell analysis, **(E)** Schematic summary of the single-cell trajectories that underlie the population response. Numbers denote the major clusters (**Fig. 3**) over time. The green trajectory denotes the drug-free baseline. Without antibiotic (centre), cultures progress through a natural growth cycle from mixed transcriptional states to stationary-like states (green arrow). The light-blue trajectory denotes the low-concentration route, where low-concentration exposure (left, 0.5 mg/L) drives an early envelope-stress programme (cluster 13; markers include *GlsB/NirD* and related stress-response loci), producing a transient detour followed by recovery to the growth trajectory. The orange trajectory denotes the high-concentration route. High-concentration exposure (right, 2 mg/L) selects an acute OM repair/efflux state (cluster 4; markers include *macAB–tolC, adeABC, lolA/tolA, bfmS, ompA*), which feeds into a slow-growing tolerant branch enriched for oxidative and translational-stress signatures. Lightning bolts denote antibiotic challenge; icons illustrate envelope damage, OM repair/efflux and stress readouts. This panel distils the early concentration-specific bifurcation observed by scRNA-seq and links it to the genetic and physiological phenotypes in panels **A–D**.

Imaging readouts at 1 h reinforce these fitness effects and provide mechanistic insights. Under 2 mg/L, mutants in envelope maintenance and efflux/trafficking (*Tn::tolA*, *Tn::lolA*, *Tn::adeA*) and in periplasmic-stress modules (*Tn::ABUW_RS07170*, *Tn::ABUS_RS18980*) displayed significantly elevated PI, membrane depolarisation, and oxidative stress compared to WT (**Fig. 4B-D**, right). These profiles indicate compromised outer-membrane integrity, loss of membrane potential and heightened oxidative burden, where biophysical liabilities expected from failure of lipoprotein delivery and MacAB–TolC/AdeABC-mediated drug extrusion. At 0.5 mg/L, the distributions of PI, DiBAC_4_(3) and CellROX fluorescence in these mutants were largely indistinguishable from the wild-type at 1 h (**Fig. 4B–D**, middle); consistent with the single-cell observation that low-concentration response is a transient, recoverable envelope-stress detour rather than a sustained injury state. As a result, strong physiological differences were not yet apparent. Collectively, the single-cell imaging phenotypes show that, under high polymyxin pressure, disruption of any of the single-cell-nominated marker genes rendered cells into a permeable, depolarised and ROS-burdened state, which explains the accompanying fitness defects observed in a subset of mutants (**Extended Data Fig. 8**).

Together, the integrated analysis resolves two early, concentration-tuned trajectories (**Fig. 4E**). At low concentration, cells follow a ‘buffer-and-recover’ route that relies on periplasmic stress buffers (NirD/YgiW/Ydel- and GlsB/YeaQ/YmgE-family factors), osmoprotection via trehalose and chaperone guided proteostasis to stabilise the envelope and constrain secondary redox. At high concentration, cells assemble an acute outer-membrane repair and efflux response programme that couples lolA-mediated lipoprotein trafficking, envelope cohesion, porin remodelling/damage sensing with MacAB-TolC and AdeABC-driven drug export, prioritising survival over growth (**Fig. 4E**). The marker genes defined the dominant subpopulations captured at the early time point; independently, their mutants showed concentration-dependent susceptibility by fitness, envelope/membrane integrity, and ROS burden, with the strongest effects at 2 mg/L. Bulk RNA-seq hinted at envelope-centric defences but could not assign them to cells. The single-cell framework identified subpopulation-restricted programs and nominated specific genes (e.g., *adeA*, *bfmS*, *macB*, *tolA*, *ABUW_RS18980*, *ABUW_RS07170*) whose perturbation demonstrably alters the outcome of polymyxin exposure. This concordance between transcriptional state, pathway function, and mutant phenotype elevates the single-cell programmes from descriptive states to actionable mechanisms that shape survival under antibiotic exposure.

## Discussion

Phenotypic heterogeneity in antibiotic responses is an emerging theme across diverse bacterial species^8, 19–21^; yet population-based approaches have so far precluded a quantitative, time- and concentration-resolved map that assigns defence programmes to individual cells within the same culture and links those states to outcome. Here, we present the first dynamic single-cell atlas that captures how bacterial population splits into divergent survival trajectories under antibiotic stress.

Polymyxin was chosen as the model antibiotic here due to its critical role as a last-line therapy worldwide^22^, providing a clinically meaningful framework for understanding bacterial stress responses. We show that a seemingly uniform bacterial culture is instead a mosaic of distinct transcriptional states whose composition depends on antibiotic concentration and time. By profiling 72,176 individual cells using scRNA-seq, we uncover discrete subpopulations that adapt distinct survival strategies. Some cells rapidly activate resistance and survival pathways and endure treatment, whereas others fail to adapt and are eliminated. This directly exposes a critical blind spot in bulk RNA-seq: while bulk analyses painted a broad response (e.g., envelope remodelling, efflux pump activation), scRNA-seq reveals how these responses are partitioned into distinct cellular subgroups following different temporal trajectories.

A central discovery of our study is the concentration-dependent bifurcation of cell fates under antibiotic treatment. Rather than a one-size-fits-all response, the population split into two major divergent trajectories. Under low-concentration treatment, 91.4% of cells transiently enter an envelope-stress state and then return to a growth-oriented path. In contrast, under high concentration, 70.1% of cells are rerouted into tolerant states with little prospect of re-entering exponential growth within the experimental timeframe. Viability measurements confirmed theses transitions: only a small subset of cells (i.e., 0.063% of initial cells) survived high-concentration treatment, yet these survivors ultimately fueled a rebound in population density. Such tolerance-driven relapse exemplifies a major clinical challenge in treating bacterial infections, whereby even a minute reservoir of surviving cells can repopulate after therapy ends^15,23,24^.

Mechanistically, the scRNA-seq data disentangle the composite stress response into discrete modules executed by distinct subpopulations. Along the high-concentration branch, surviving cells prioritised envelope-maintenance, lipoprotein trafficking, phospholipid transport, MacAB-Tol system-mediated efflux, proteostasis and ribosome rescue, consistent with a membrane-first injury that propagates to protein homeostasis. Concurrently, other cells preferentially activated antioxidant defenses (reflecting secondary redox stress) or engaged trans-translation and associated factors, marking translational stalling. Under low-concentration treatment, a coordinated but short-lived envelope-stress programme dominated early and then receded, allowing most cells absorbed a modest hit and resumed the normal growth trajectory over the following hours. Together, these distinct patterns show that envelop repair, transport and stress-mitigation modules are not uniformly co-activated in every cell but are distributed across specialised subpopulations. Importantly, this heterogeneity is not a static trait of the culture. Instead, it is structured, time-resolved and dynamically tuned into antibiotic intensity, with differing modules taking the lead depending on stress severity.

Critically, the single-cell map of transcription and trajectory is anchored in function. Perturbing genes that mark the dominant early programmes altered 24-h fitness relative to wild type and, at the high concentration, produced rapid increases in envelope permeability, membrane depolarisation and reactive-oxygen burden within one hour. These convergent genetic and physiological effects indicate that the subpopulation programmes identified at single-cell resolution encode liabilities that shape survival. Importantly, the phenotype was not strictly cluster-specific: genes originating from the marker sets of both the low- and high-concentration groups contributed to survival under membrane-targeting stress. We therefore interpret these transcriptional programmes as an experimentally validated tolerance architecture that highlights envelope maintenance, lipoprotein trafficking and efflux, proteostasis and redox control as tractable targets for interference.

These mechanistic insights gained in the present study offer important clinical implications for improving the precision and effectiveness of antibiotic treatment in patients. We deliver the first high-resolution, longitudinal map of population structure, a level of temporal detail not previously attained in the antimicrobial pharmacology field. This rich temporal resolution allowed us not only to identify distinct cell subpopulations, but importantly, to elucidate their dynamics of emergence and decline over the course of treatment. The ability to track subpopulations dynamics is critical for understanding how to optimise antibiotic use. For example, low-concentration polymyxin exposure (cluster 13 at 1 h) should be minimised, as those cells initially showing envelope-stress response had largely recovered after 4-7 hours and reverted to a growth-oriented trajectory. In contrast, sufficiently high concentrations of polymyxin should be achieved to rapidly eradicate the survivors in a tolerant-like dormancy. If polymyxin monotherapy is not possible to eliminate tolerant state cells, pairing polymyxin with another agent should be considered to target the specific tolerance mechanisms we identified. Examples could include inhibitors of lipid A modification (to prevent polymyxin resistance via envelope remodelling), or ROS-generating compounds that overwhelm the bacterial antioxidant defenses^25–29^. Such adjunctive strategies could eradicate the last surviving cells that would otherwise sustain chronic or relapsing infection. More broadly, by uncovering distinct evolutionary trajectories previously hidden in bulk RNA-seq, our single-cell framework reveals a critical vulnerability in current antibiotic dosing strategies and demonstrates that precision therapy is essential to prevent the evolutionary emergence of resistance.

This study is not without limitations, which in turn define promising avenues for future exploration. As our analyses were conducted *in vitro*, extending this framework to biofilm systems and in vivo contexts will be important for resolving the complete state space and its context-dependent transitions. Another limitation is that this proof-of-concept study focuses on a subset of markers for functional validation; perturbations within each module and the combinations of multiple modules, alongside antibiotic treatment, will be critical to identify novel combinations that collapse survival trajectories. Such studies will be important for validating potential drug targets and combination therapies suggested by our scRNA-seq analyses.

Overall, we map a dynamic, concentration-structured landscape of antibiotic responses at the single-cell level, revealing diverse survival trajectories that are obscured in conventional population-averaged analyses. By integrating both bulk and single-cell transcriptomics, we resolve an early bifurcation in bacterial fate and identify the modules that dominate each branch, with functional perturbation linking these modules causally to survival. These single-cell findings strongly support mechanism-guided precision therapy, particularly by employing novel combinations to intercept cultures while they traverse transient, susceptible states. Importantly, the framework is readily transferable across other antibiotics, bacterial species and clinical settings. By turning population averages into actionable single-cell mechanisms, our single-cell approach charts a path toward precision therapy to improve treatment outcomes in patients and minimise emergence of multi-drug resistance.

## Online Methods

### Bacterial strain and culture conditions

We used *Acinetobacter baumannii* strain AB5075 (a modern model strain for virulence and antibiotic resistance studies) for all experiments^30^. Bacteria were grown in cation-adjusted Mueller-Hinton broth (CAMHB) at 37 °C with shaking (200 rpm), unless otherwise noted. For routine cultivation, overnight cultures were started from a single colony and grown to stationary phase (~12 h). Prior to the experiments, an overnight culture was diluted 1:100 into fresh CAMHB and grown to mid-log phase (OD_600nm_ = 0.5) to ensure a predominantly dividing, planktonic population. This mid-log culture was used as the starting inoculum for all conditions.

### Antibiotic treatment, sampling and bright field imaging

Polymyxin B sulphate (Sigma-Aldrich) was prepared as a stock solution (1 mg/mL in water). At time 0 h, polymyxin B was added to aliquots of the log-phase *A. baumannii* culture to achieve the desired concentrations: a low minimal inhibitory concentration of 0.5 mg/L and a high concentration of 2 mg/L (approximately equal to 4 times MIC for this strain, as determined by broth microdilution). A control culture received no antibiotic. All cultures (including control and treatments) started at the same cell density (~10^8^ CFU/mL) and were in the same volume of medium, to ensure comparable growth conditions aside from the antibiotic. Samples were taken at 0 (immediately before adding antibiotic), 1, 4, and 7 h after polymyxin addition to capture the early response (1 h), an intermediate response (4 h), and later-stage survivors (7 h). At each time point, an aliquot of culture was removed for analysis. A portion of each sample was serially diluted and plated on nutrient agar to enumerate viable CFUs, confirming the antibiotic’s killing effect over time as previous described^31^. In the 2 mg/L condition, viability dropped by ~99% at 1 h, then slightly increased by 7 h (consistent with regrowth of a tolerant subpopulation), whereas in the 0.5 mg/L condition, there was a smaller growth delay by 1-4 h. These viability dynamics informed our interpretation of the single-cell data and indicated successful exposure to the intended polymyxin stress (**Extended Data Fig. 6**).

For bright field imaging, parallel aliquots from the same time points (control, 0.5 mg/L, 2 mg/L at 0, 1, 4 and 7 h) were spotted (0.3 μL) onto agarose gel pads prepared according to a modified published protocol^32^ (pad composition and preparation in Supplementary Methods). Preparations were imaged immediately on a Leica DMi8 wide-field microscope equipped with a 100× oil-immersion objective (NA 1.32) under identical illumination and exposure settings within each experiment. Acquisition parameters and image-processing steps are detailed in the Supplementary Methods.

### Bulk RNA-seq library preparation and analysis

In parallel with single-cell sampling, we prepared samples for bulk RNA sequencing from the 1 h and 4 h time points of each condition (0.5 mg/L, 2 mg/L, and control), following previously described methods^33^. In brief, for each selected condition/time, approximately 10^8^ CFU/mL bacteria were pelleted and immediately stabilised using RNAprotect Bacteria Reagent (Qiagen) to prevent RNA degradation. Total RNA was extracted using the RNeasy Mini Kit (Qiagen) with on-column DNase digestion. Ribosomal RNA was removed from the samples using a Ribo-Zero Plus kit (Illumina) targeting bacterial rRNA. The remaining mRNA-enriched RNA was used to construct sequencing libraries with an Illumina TruSeq Stranded mRNA Library Prep kit (modified for no-poly(A) bacterial RNA). Libraries were indexed and sequenced on an Illumina platform (NovaSeq 6000) to generate 150 bp paired-end reads. Three biological replicates were included for each condition. Raw reads were quality-checked and trimmed for adapters with trim-galore^34^. Filtered reads were aligned to the *A. baumannii* AB5075 reference genome using subreadR^35^. Gene expression counts were obtained with featureCounts, and differential expression analysis was performed using quasi-likelihood estimation via edgeR in R^35,36^. Genes with an adjusted p-value (BH-adjusted) < 0.05 and absolute log2-fold change > 1 were considered significantly differentially expressed. Up- and down-regulated gene sets for each comparison (e.g., 1 h 0.5 mg/L vs 1 h control, 1 h 2 mg/L vs 1 h control, etc.) were subjected to functional enrichment analysis. KEGG pathway enrichment was conducted using the ClusterProfiler package in R, with pathways having p-value < 0.05 considered enriched^37^. Principal component analysis (PCA) and hierarchical clustering were performed on variance-stabilised expression data to visualise global transcriptomic differences between samples using R.

### Single-cell RNA-seq

We employed the commercially smRandom-seq platform (M20 Genomics, Hangzhou, China) for single-cell transcriptomic profiling of bacteria^38^. At each sampling time point (0, 1, 4, 7 h for each condition), approximately 10 mL of culture (~10^7^ cells) was immediately fixed by adding freshly prepared paraformaldehyde to a final concentration of 4% and incubating for 30 min at room temperature. Fixation preserves the RNA in each cell by crosslinking and halts further gene expression changes. Fixed cells were washed twice with phosphate-buffered saline (PBS) to remove residual media and polymyxin B. Cell suspensions were first counted using a Moxi cell counter and diluted according to the manufacturer’s instructions to obtain single-cell suspensions. The bacterial single-cell RNA-seq library was prepared using the VITAPilote kit (M20 Genomics). *In situ* reverse transcription was performed with random primers, and adaptor sequences were subsequently added to the resulting cDNA fragments. Droplet barcoding of individual bacteria was performed on the VITACruiser Single Cell Partitioning System (M20 Genomics, Hangzhou, China), in which bacteria, DNA extension reaction mix, and hydrogel barcoded beads were co-encapsulated. Following droplet generation, the aqueous phase containing cDNAs was purified with magnetic beads, amplified by PCR, and subjected to a second purification step. The resulting products were pooled to construct sequencing-ready libraries.

### Preprocessing and clustering of single-cell data

Sequencing was done on a PE150 (Illumina) and raw sequencing reads from the single-cell run were processed with a custom pipeline. First, we extracted cell barcodes and unique molecular identifiers (UMIs) from Read 1, and trimmed adapters from Read 2. We mapped Read 2 sequences (cDNA inserts) to the *A. baumannii* AB5075 reference genome using BWA-MEM^39^. Only reads that mapped uniquely to gene features were retained. We then collapsed reads into UMIs to obtain digital expression counts for each gene in each cell. The initial digital expression matrix (genes × cells) was filtered to remove low-quality cells and likely droplets containing no cell or multiple cells. Specifically, we excluded barcodes with fewer than 50 genes detected or with an extraordinarily high gene count (>3 standard deviations above the mean, indicative of multiplets). After filtering, we normalised the data by library size (UMI counts per cell) and log-transformed the counts.

For clustering, we used the Seurat R package (v5.0)^40^. We identified highly variable genes across the dataset and performed PCA on the centred, scaled expression values of these genes. Significant principal components were determined by examining the PCA elbow plot and JackStraw test. We then constructed a *k*-nearest-neighbor graph in the PCA space (using *k*=20) and applied the Louvain algorithm for graph community detection (with resolution parameter 0.6) to define clusters of cells. This yielded 15 clusters.

Marker genes for each cluster were identified using Seurat’s *FindMarkers* function (Wilcoxon rank-sum test), comparing each cluster to all others. We applied Bonferroni correction for multiple testing. Genes with an adjusted p-value < 0.01 and at least 0.25 log_2_-fold change were considered significantly enriched in a given cluster. These marker genes were used to infer the biological functions of each cluster, as described in the Results. We visualised the expression of key marker genes on the FDL plot and with heatmaps/dot plots to confirm that their patterns corresponded with cluster assignments^41^. To quantify cluster composition across samples, we calculated the proportion of cells from each condition/time that fell into each cluster.

### Pseudotime trajectory inference and manifold dynamics analysis

To reconstruct potential lineage relationships between the identified clusters/states under polymyxin exposure, we combined graph-based lineage inference with manifold-level density analysis. We first performed pseudotime trajectory reconstruction using the Scanpy toolkit (v1.9) in Python, which includes the partition-based graph abstraction (PAGA) method^42,43^. We converted our Seurat-processed data to AnnData format for Scanpy, using the log-normalised expression values and the cluster labels determined in Seurat as cell annotations. We first constructed a *k*-nearest-neighbour graph of cells (using *k*=15 neighbours) with Scanpy’s *pp.neighbors* function on the PCA-reduced space. We then applied *tl.paga* to compute the PAGA connectivity between clusters. PAGA treats each cluster as a node and estimates connections (edges) based on shared nearest-neighbours between cells of different clusters, yielding a graph where edge weights represent the confidence of transitional relationships between clusters.

The PAGA graph was visualised using a force-directed layout (**Fig. 3F**), with each node labelled by cluster identity and additionally annotated post hoc with the predominant condition/time it represented (for clarity in interpretation). PAGA revealed a branching topology among the clusters, as described in the Results. We further explored pseudotemporal ordering of cells by choosing an appropriate start and end for diffusion pseudotime (DPT) analysis. Based on the experimental design, we set the 0 h baseline cluster (cluster 5, one of the stationary-phase starting clusters) as the root of the trajectory, and we treated cluster 3 (adaptive resistance persister state) and cluster 4 (acute stress response state) as two potential end-points of interest. Using Scanpy’s *tl.dpt* function, we computed a pseudotime value for each cell, which orders cells along a trajectory taking into account the PAGA connectivity (this effectively spreads out cells along the graph respecting the branch structure).

To complement the PAGA-derived connectivity map with a geometric representation of how cell populations redistribute through transcriptional space, we performed a manifold-based density and flow analysis. All real single-cell transcriptomes from the 0, 0.5 and 2 mg/L conditions (0–7 h) were embedded into a shared FDL), providing a low-dimensional manifold that preserves global transcriptomic geometry. For each concentration-specific trajectory, we defined a common baseline using the untreated control, 0 h population and combined these cells with those from subsequent timepoints (1, 4, 7 h) of the same condition. Two-dimensional Gaussian kernel density estimates (KDEs) were computed on the FDL coordinates to quantify how cells occupy the manifold under each condition^42^. Density values were normalised within each concentration and rescaled to a shared 0.2 – 1.0 range, enabling direct comparison across conditions. For visualisation, densities were plotted using a continuous Spectral-derived colormap (blue → yellow → red), capturing shifts from low- to high-density regions. To illustrate directional population flow, we computed the centroid of the cell distribution at each timepoint and connected these centroids with arrows, representing the dominant transcriptomic trajectory from baseline through 1, 4 and 7 h. Together, the PAGA-derived lineage structure and the manifold-level density trajectories provide a unified view of how polymyxin exposure reshapes transcriptional state space, revealing both the branching architecture of stress responses and the geometric flow of cells through the underlying manifold.

### Kinetic growth and fitness assay

For 24 h growth curves, we used a Cerillo Microplate Reader System with temperature control and continuous shaking. Mid-log cultures were diluted into assay medium to a uniform starting inoculum and dispensed into clear, flat-bottom 96-well plates alongside medium blanks and wild-type (WT) controls on every plate and for every concentration. Optical density at 600 nm (OD_600_) was recorded kinetically for 24 h at fixed intervals (every 10 mins). For each well, OD_600_ time series were blank-subtracted and analysed in R (v4.3.2) using the *growthcurver* package to estimate logistic parameters and the area under the growth curve (AUC; 0–24 h)^44^. Fitness was summarised as WT-normalised log_2_AUC:

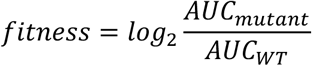

Values were computed within plate and concentration to control for run-to-run variation, then averaged across biological replicates for the heat map (**Fig. 4A**). Representative 24 h trajectories for selected mutants are shown in **Extended Data Fig. 7**. Wells failing prespecified QC (irregular baselines, condensation artefacts, or non-monotonic early traces) were excluded prior to fitting.

### Fluorescence imaging of AB5075 transposon mutants

Transposon insertion mutants of *A. baumannii* AB5075 were selected from the single-cell RNA-seq marker sets that dominate at 1 h (**Fig. 2D**) and imaged alongside wild-type controls. Mid-log cultures were split and exposed for 1 h to 0.5 mg/L or 2 mg/L polymyxin B, or left untreated, using matched inocula and volumes as in the main assays; cells were then stained at room temperature for 10 min with DiBAC₄(3) (10 µM), propidium iodide (PI; 2.5 mg/L) and CellROX™ Deep Red (2.5 µM) in the dark. A 0.3 µL aliquot of each sample was mounted on agarose gel pads prepared according previously reported protocol^32^ and imaged immediately on a Leica DMi8 wide-field microscope with a 100× oil-immersion objective (NA 1.32). Four channels were acquired per field: bright-field (transmission) for morphology, FITC (Ex 460–500 nm, Em 512–542 nm) for DiBAC, RHOD (Ex 541–551 nm, Em 565–605 nm) for PI, and Y5 (Ex 590–650 nm, Em 662–738 nm) for CellROX; illumination and exposure were held constant across strains within each concentration, and multiple fields were collected across ≥3 independent experiments. Single-cell segmentation used Cellpose^45^ with morphology filters to exclude debris/aggregates; per-cell intensities were background-subtracted, log₁₀-transformed, and normalised to the WT median acquired in the same batch and concentration. Summary distributions (**Fig. 4B–D**) and statistical tests are detailed in the Statistics and Supplementary Methods sections.

### Statistics and data visualisation

All the analyses were performed in R without further clarification. For growth curves, *growthcurver* returned AUC and logistic parameters; fitness values were aggregated as means ± s.e.m. per mutant/condition. For imaging, per-cell WT-normalised intensities were modelled with linear mixed-effects frameworks (mutant as fixed effect; batch and FOV as random intercepts) to account for imaging heterogeneity; pairwise comparisons to WT used estimated marginal means with Benjamini–Hochberg correction across mutants. Heat maps and violins plots were generated with *ggplot2*; colour scales and panel layouts were fixed across concentrations.

## Data availability, reproducibility and supplemental information

Processed RNA-seq count matrices are available via Figshare (https://doi.org/10.6084/m9.figshare.30745412). Supplementary data are available at https://osf.io/r5j8e/files/xm8yr. Code used for data processing, analysis and figure generation is available on GitHub (JinxinMonash/scRNA-seq-in-Acinetobacter-baumannii).

## Acknowledgements

J.L. is an Australian National Health Medical Research Council (NHMRC) Investigator Research Fellow (APP2025937) and T.V. is an Australian Research Council (ARC) Industrial Fellow (IM230100534). Y.W.L. is currently also an employee of Certara. This work was partially supported by grants from Centre of Excellence for Antimicrobial Therapeutics Discovery and Innovation (CEATDI), Monash Suzhou Research Institute (MSRI8002003) and Consortium for Infection and Innovation (CII), Monash University - Wenzhou Medical University Biomedical Alliance, The First Affiliated Hospital of Wenzhou Medical University (2025WMU-X001).

## Notes

https://doi.org/10.6084/m9.figshare.30745412

https://osf.io/r5j8e/files/xm8yr

### Competing Interest Statement

The authors have declared no competing interest.

